# Focal brain stimulation sites that modify autonomic arousal map to a convergent brain circuit and potential therapeutic target

**DOI:** 10.64898/2026.01.08.698446

**Authors:** Ryan D. Webler, Clemens Neudorfer, Eva S. A. Dijkstra, Summer B. Frandsen, Nicole Chiulli, John Miller, Ru Kong, Sanaz Khosravani, Garance M. Meyer, Alexandre Boutet, Jurgen Germann, Gavin J.B. Elias, Nicholas L. Balderston, Andrew Strohman, Wynn Legon, Jordan H. Grafman, Martijn Arns, Andres M. Lozano, Shan H. Siddiqi

## Abstract

Causal modulation of autonomic outflow could yield new therapeutic targets for autonomic hyperactivation. We employed three natural experiments in which different brain regions were targeted using transcranial magnetic stimulation (TMS) (n=139 sites, n=14 individuals), deep brain stimulation (n=392 sites, n=58 individuals), or low-intensity focused ultrasound (n=46 sites, n=23 individuals) with subsequent autonomic measurements. Using a human connectome database (n=1000) as a wiring diagram, we identified a convergent brain circuit that, when focally modulated, transiently reduces autonomic arousal. This circuit significantly resembled previously reported causal circuits for posttraumatic stress disorder (PTSD) and anxiety. In independent datasets, TMS to the autonomic arousal circuit reduced laboratory startle in healthy volunteers (n=28), lesions to this circuit reduced exaggerated startle in PTSD (n=193), and TMS to this circuit reduced anxiety-related autonomic symptoms in patients with clinically significant anxiety (n=30). Thus, the convergent circuit may serve as a potential neuromodulation target for autonomic hyperactivation.

---

Autonomic arousal is a coordinated set of physiological adjustments, such as increased heart rate and blood pressure, that support adaptive responses to environmental demands (1). While transient autonomic activation is an appropriate response to challenge, excessive activation can cause significant distress and impairment (2). Hyperactivation to anxiogenic stimuli is a core feature of anxiety and trauma-related disorders (3,4). Similar patterns of dysregulated autonomic activity also occur across neurological, cardiac, endocrine, and other conditions (5–7). Although symptom-specific treatments such as beta-adrenergic blockade can be effective in some cases, broader interventions for autonomic hyperactivation remain limited.

Focal neuromodulation methods including transcranial magnetic stimulation (TMS), deep brain stimulation (DBS), and low-intensity focused ultrasound (LIFU) can modulate activity in specific human brain circuits (8–10). These interventions have been shown to be therapeutic for some neuropsychiatric and neurological indications. However, their focality requires that the right location be targeted to treat a given indication. Causal circuit mapping is an emerging method that can be used to identify circuit-based neuromodulation targets for specific indications (11). This technique uses brain maps (12) to trace common circuits connecting spatially disparate causal interventions, such as brain lesions or brain stimulation targets that selectively modify the same outcome. Prospective stimulation of causally-derived circuitry for a given outcome has been shown to modify that outcome (13–16)

Building on these findings, the present study used causal circuit mapping to derive a novel circuit that might serve as a potential neuromodulation target for autonomic hyperactivation. First, we derived independent circuits connecting TMS (n = 14, stimulation sites = 139), DBS (n = 58, stimulation sites = 392), and LIFU (n = 23, stimulation sites = 46) sites that transiently modulated different autonomic outcomes. Second, we established alignment between these circuits and combined them into a convergent autonomic circuit. Third, we performed post hoc analyses to examine relationships between this convergent circuit and sustained changes in anxiety-related autonomic activation across multiple independent datasets.

## Results

### TMS circuit associated with greater heart rate deceleration

The TMS dataset includes 14 healthy subjects who received stimulation to 10 different cortical targets (17) (Figure 1a). Targets were selected based on individualized functional connectivity to the subgenual anterior cingulate cortex (sgACC), as previously reported (17). This target selection procedure was motivated by the hypothesis that indirect modulation of the sgACC partially mediates the antidepressant and autonomic effects of TMS (18). Targets were stimulated using a validated protocol shown to yield temporary heart rate deceleration entrained to TMS. This protocol includes 5-second trains of 10 Hz TMS, which briefly decelerate heart rate, separated by an 11-second inter-trial interval during which heart rate normalizes. TMS induced entrainment of heart rate deceleration at each target was the primary study outcome. One target was removed due to data corruption, leaving a total of 139 target sites.

**Figure 1:**
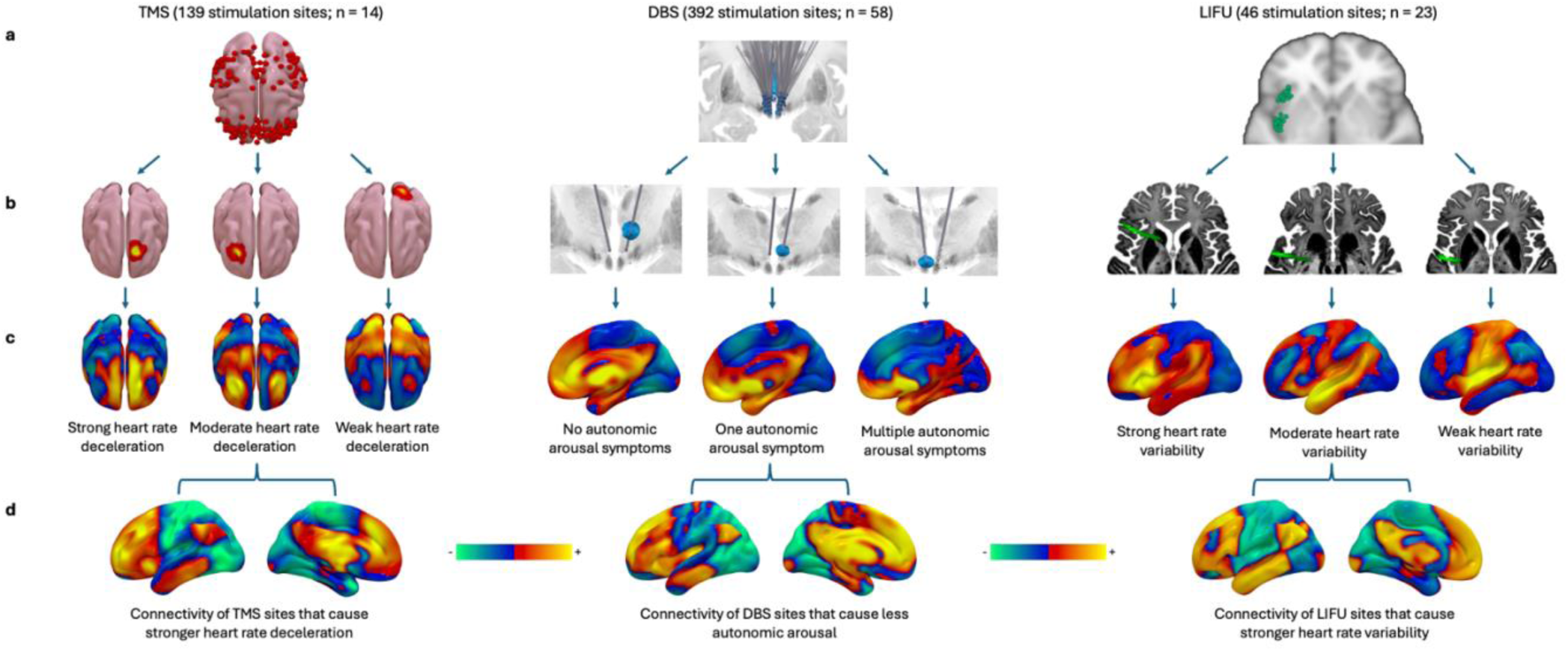
Generating TMS, DBS, and LIFU autonomic arousal circuits. a. Spatially variable TMS (n =139), DBS (58 leads, 392 sites), and LIFU (n = 46) target locations. b. TMS sites were estimated using concentric spheres of decaying intensity (up to 12mm); DBS sites were estimated using virtual tissue activation modeling; LIFU sites were estimated using acoustic modeling. c. Functional connectivity profiles of brain stimulation sites were mapped using a normative connectome. Connectivity profiles associated with different levels of autonomic arousal are depicted. d. Voxel-wise TMS, DBS, and LIFU maps were generated using repeated measures linear mixed-effects models. These models tested relationships between functional connectivity and autonomic arousal (controlling for within-subject effects) at every voxel. Warm and cool colors reflect positive and negative connections of brain stimulation sites associated with less autonomic arousal, respectively.

To map a TMS heart rate deceleration circuit, we modeled TMS sites as concentric spheres of decaying intensity (up to 12mm) as in prior work (19) (Figure 2b), mapped the functional connectivity of each TMS site using a normative connectome database (n = 1,000) (12) (Figure 2c), and used linear regression to assess relationships between TMS site connectivity and heart rate deceleration at each voxel (Figure 2d). A repeated measures mixed-effects model was used to test between-subject effects while accounting for within-subject correlations resulting from multiple TMS sites per subject. This resulted in a functional circuit comprised of TMS site connections associated with stronger or weaker heart rate deceleration.

**Figure 2:**
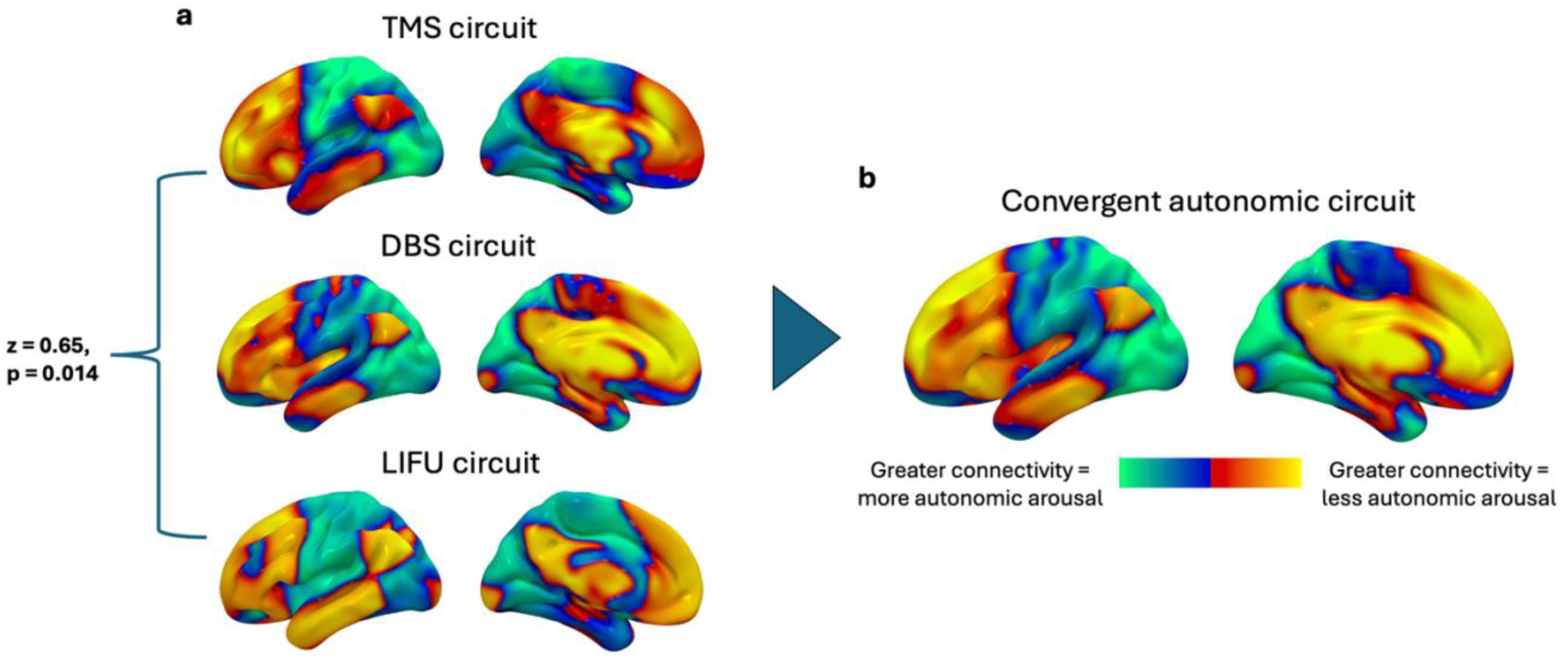
Convergent autonomic circuit. a. Pooled pairwise voxel-wise Pearson correlation was performed to assess similarity between the TMS, DBS, and LIFU circuits; permutation methods were used to test significance. The circuits were more similar than expected by chance. b. TMS, DBS, and LIFU circuits were combined into a weighted mean convergent autonomic circuit. Warm and cool colors reflect positive and negative connections of brain stimulation sites associated with less autonomic arousal.

Highly similar circuits were generated using electric field (E-field) modeling to estimate TMS stimulation volume across multiple thresholds (mean r = 0.89), but these were not treated as primary analyses because normative connectome analysis has not been independently validated with E-fields (see Supplementary Materials).

### DBS circuit associated with less autonomic arousal

The DBS dataset includes 58 patients with mild Alzheimer’s disease who received acute stimulation to hypothalamic area targets (392 total targets) (20) (Figure 1a). This dataset was chosen because it captures a wide range of DBS-related autonomic symptoms including cardiovascular (tachycardia, hypertension), thermoregulatory (warmth, flushing, sweating, coldness), gastrointestinal (nausea), and vestibular (dizziness) changes.

To map a DBS autonomic arousal circuit, we first estimated DBS site locations using volume of tissue activation (VTA) modeling, as previously reported (21) (Figure 1b). Next, we mapped the functional connectivity of each DBS site using the same normative connectome (12) used to derive the TMS circuit (Figure 1c). As with the TMS circuit, we employed repeated measures linear mixed-effects models to assess the relationship between DBS site connectivity and autonomic arousal at each voxel (Figure 1d).

To ensure that results were not biased by analytical choices, we repeated the analysis by: (1) summing individual autonomic symptoms (rather than summing across autonomic categories, i.e., cardiovascular, thermoregulatory, gastrointestinal, vestibular) (correlation to primary circuit: r = 0.99), (2) generating circuits from individual symptoms and then averaging them (correlation to primary circuit: r = 0.98), and (3) generating averaged circuits after leaving one symptom out (mean correlation to primary circuit: r = 0.98) (see Supplementary Materials).

### LIFU circuit associated with greater heart rate variability

The LIFU dataset includes 23 healthy participants who received modulation to individualized anterior and posterior insula targets (46 total targets) identified using an insular atlas (22) (Figure 1a). One second pulse trains of 500 kHz (1 kHz pulse-repetition frequency, 36% duty cycle) LIFU were delivered to insular targets using a single-element focused transducer (Sonic Concepts H-281). This dataset was chosen because heart rate variability, a cardiac function mediated by the autonomic nervous system (23), was a primary study outcome.

To map a LIFU heart rate variability circuit, we modeled LIFU sites using acoustic modeling as described in Legon et al. (2018) (24) (Figure 1b). Stimulation sites were thresholded at >50% peak energy (10). Next, we mapped the functional connectivity of LIFU sites using the same normative connectome used to derive the TMS and DBS circuits (Figure 1c). We then used repeated measures linear mixed-effects models to test the relationship between LIFU site connectivity and heart rate variability at each voxel (Figure 1d).

As with the TMS and DBS circuits, we generated multiple iterations of the LIFU circuit to ensure that our result was not driven by analytical choices. LIFU circuits generated using acoustic thresholds of 25% (r = 0.90) and 75% (r = 0.98) were highly similar to the primary LIFU circuit thresholded at 50% (see Supplementary Materials).

### Convergent autonomic circuit

To assess for convergence between the TMS, DBS, and LIFU circuits, we performed pairwise spatial Pearson correlations and averaged the Fisher z-transformed results. To assess significance, we generated a null distribution for each dataset using a permutation method that randomly paired autonomic outcomes from one stimulation site with another site, similar to our prior work (13). The average pairwise circuit correlation was stronger than expected by chance (mean z = 0.65, p = 0.014) (Figure 2a), suggesting shared autonomic circuitry across all three stimulation modalities.

Given significant convergence between the TMS, DBS, and LIFU circuits, we combined them into a weighted mean autonomic arousal circuit (Figure 2b). We corrected for voxel-wise multiple comparisons using the permutation-based tStat method, as per consensus guidelines (25). After multiple comparisons correction, the convergent autonomic circuit contains significant positive connections in the dorsomedial prefrontal cortex, ventrolateral prefrontal cortex, anterior cingulate cortex, anterior insula, thalamus, posterior cingulate cortex, middle temporal gyrus, midbrain, and cerebellum. Significant negative connections are in the paraventricular hypothalamus, infundibulum, inferior parietal lobule, temporal pole, and occipital cortex (see Supplementary Materials). Significant peaks include the fornix, a fiber tract that projects to the hypothalamus (26), and the anterior midcingulate cortex, a location found to mediate autonomic responses like tachycardia and heat sensations in previous neuromodulation studies (27–30).

### Relevance to anxiety-related autonomic activation

In a series of post hoc analyses, we probed the convergent autonomic circuit’s relevance to anxiety-related autonomic activation in a laboratory task and in clinical patients. First, we tested whether incidental intersection of TMS sites with the convergent autonomic circuit is associated with anxiety-related activation to a laboratory anxiety task. Second, we tested: (1) whether the convergent circuit is aligned with causal anxiety-related circuits (14,31), and (2) whether incidental intersection of lesions and TMS sites with the convergent autonomic circuit is associated with sustained changes in anxiety-related hyperactivation symptoms. As in previous analyses, statistical significance was assessed using permutation testing.

#### Laboratory anxiety-related activation

Data came from a double-blinded, randomized, cross-over trial in which 28 healthy participants received active intermittent theta-burst stimulation and sham stimulation (order randomized) to the right dorsolateral prefrontal cortex 24 hours before a laboratory anxiety task (Figure 3a) (32). Anxiety-related activation was measured using anxiety potentiated startle (APS), which indexes startle magnitude changes potentiated by anticipation of an uncertain threat. We mapped TMS site connectivity using the same procedure as the heart rate deceleration analysis above (12), and correlated TMS-site seed-maps with the convergent autonomic circuit. Active TMS sites with stronger connectivity to the convergent autonomic circuit reduced anxious arousal more than sham (t = 2.72, permuted p = 0.03) (Figure 3b).

**Figure 3:**
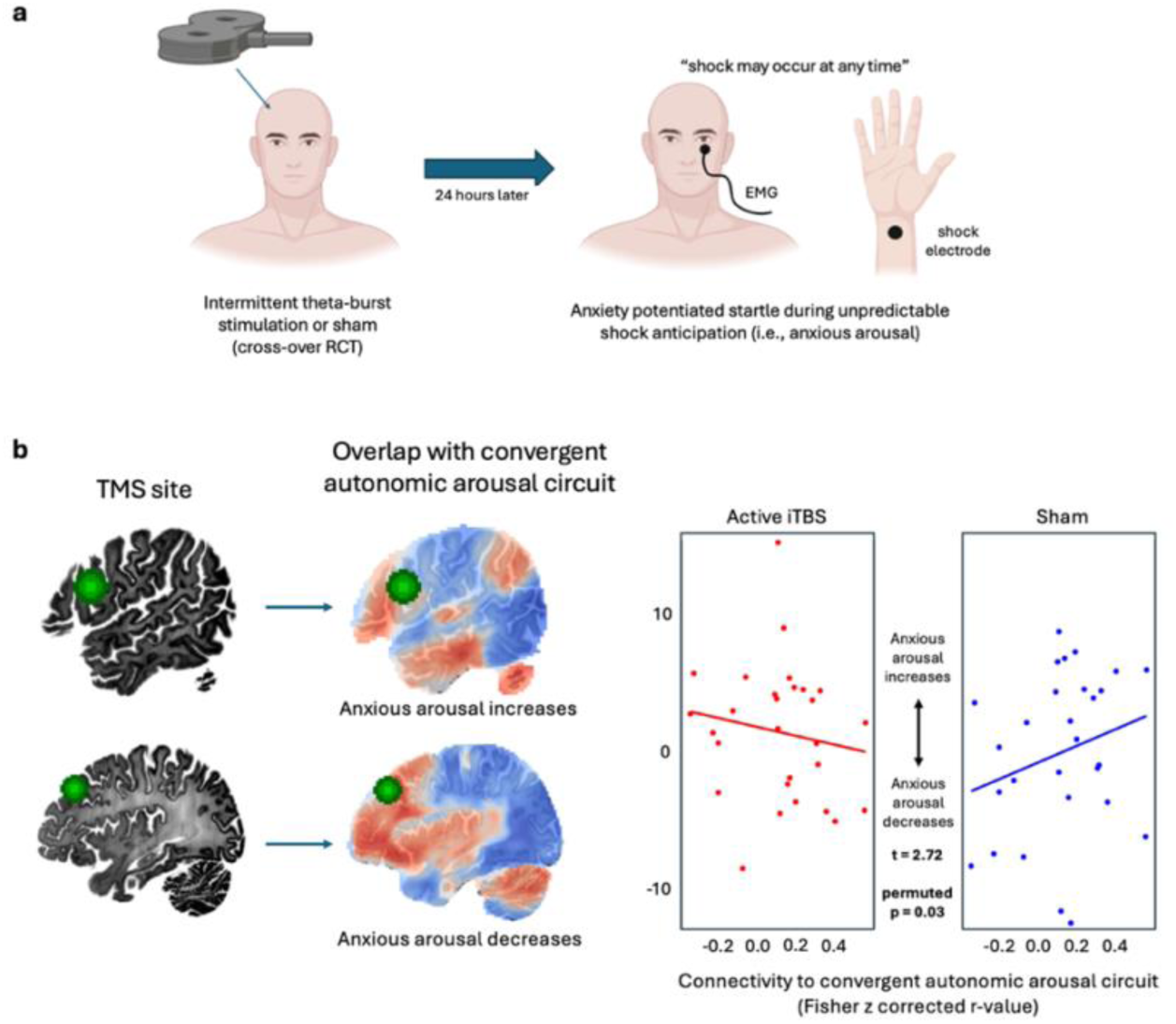
Relevance of the convergent autonomic circuit to laboratory anxious arousal. a. Data from a double-blinded, randomized, cross-over trial in which 28 healthy participants received intermittent theta-burst or sham to individualized right dorsolateral prefrontal cortex targets. One day after TMS, participants underwent a laboratory anxiety task. Anxiety-related activation was measured using anxiety potentiated startle which indexes the difference in startle response magnitude during anticipation of an unpredictable mild electrical shock versus baseline. b. Intersection of example TMS sites with the convergent autonomic circuit are depicted. Active TMS sites with stronger positive connectivity to the convergent autonomic circuit reduced laboratory activation more than sham.

#### Clinical anxiety-related hyperactivation

PTSD data came from Phase 3 of the Vietnam Head Injury Study (VHIS) (n = 193) (33), in which a proportion of Vietnam War Veterans developed PTSD (n = 61) following penetrating brain trauma. From this data, a recently published PTSD circuit (31) was derived by correlating the connectivity of penetrating head injury lesions with PTSD status (i.e., no PTSD, subthreshold PTSD, PTSD) at every voxel, independent of depression severity and lesion size. This yielded a map in which warm and cool colors reflect positive and negative functional connections of PTSD-protective lesions, respectively (Figure 4a).

**Figure 4:**
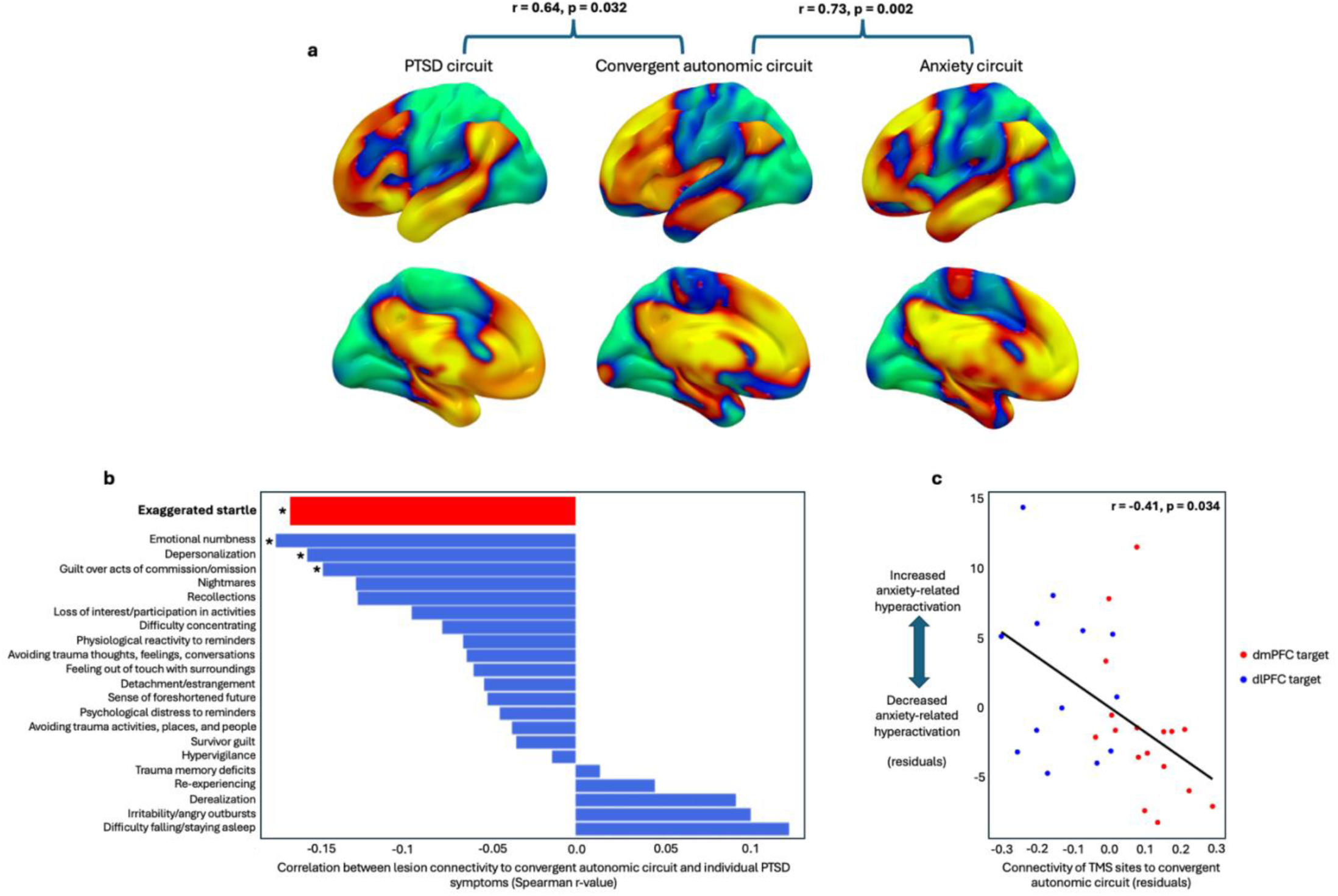
Relevance of the convergent autonomic circuit to clinical anxious arousal. a. Causal PTSD and anxiety circuits are more similar to the convergent autonomic circuit than expected by chance. b. Lesion connectivity to the convergent autonomic circuit is significantly negatively associated with specific PTSD symptoms including exaggerated startle, emotional numbness, depersonalization, and guilt over acts of omission/commission. Asterisks denote p<0.05. c. TMS site connectivity to the convergent autonomic circuit is significantly negatively associated with anxiety-related hyperactivation symptoms measured using the Beck Anxiety Inventory. Dorsolateral prefrontal cortex (dlPFC) and dorsomedial prefrontal cortex (dmPFC) target data are depicted in blue and red, respectively.

Spatial Pearson correlation was used to test alignment between the PTSD and convergent autonomic circuits. Permutation testing confirmed that the circuits are more similar than expected by chance (r = 0.64, p = 0.032). In additional analyses, we tested relationships between the convergent autonomic circuit and specific PTSD symptoms using Spearman correlation.

Lesion connectivity to the convergent autonomic circuit was negatively associated with exaggerated startle (r = -0.17, p = 0.022) – an anxiety-related hyperactivation symptom – and emotional numbness (r = -0.17, p = 0.017), depersonalization (r = -0.16, p = 0.033), and guilt over acts of omission/commission (r = -0.15, p = 0.045) (Figure 5b).

Clinical anxiety data came from multiple brain lesion (k = 2, n = 451) and brain stimulation (k = 2, n = 111) datasets that included an anxiety measure. The VHIS study used to derive the PTSD circuit was one of the lesion datasets; 181 Veterans with anxiety ratings were included. Despite heterogeneity in causal methods, study samples, and anxiety measures, anxiety (independent of depression) mapped to a common circuit across datasets (14). As with the PTSD circuit, the convergent autonomic circuit is more similar to the anxiety circuit than expected by chance (r = 0.73, p = 0.002). In additional analyses, we used data from a recent randomized comparative TMS trial (n = 30) to test whether incidental TMS site connectivity to the convergent autonomic circuit predicts changes in Beck Anxiety Inventory (BAI) hyperactivation symptoms, independent of depression. This trial compared the efficacy of dorsolateral and dorsomedial prefrontal cortex targets in patients with moderate to severe depression and anxiety. As expected, TMS sites that were more strongly connected to the convergent autonomic circuit yielded greater reductions in anxiety-related hyperactivation symptoms, independent of depression (Spearman r = -0.41, p = 0.034).

#### Depression

The depression circuit was derived from the same dataset as the PTSD circuit by correlating lesion connectivity with Beck Depression Inventory (BDI) scores at each voxel. Unlike the PTSD and anxiety circuits, the depression circuit does not align with the convergent autonomic circuit (r = 0.09, p = 0.61).

## Discussion

We derived distinct brain circuits connecting TMS, DBS, and LIFU sites that transiently reduced autonomic arousal. These circuits are more similar than expected by chance. A convergent autonomic circuit formed by combining these circuits was associated with reductions in laboratory and clinical anxiety-related autonomic activation across multiple independent datasets. Specifically, TMS sites that incidentally intersected the convergent autonomic circuit reduced laboratory anxiety-potentiated startle. Similarly, lesions and TMS sites that incidentally intersected the convergent autonomic circuit yielded sustained reductions in PTSD-related exaggerated startle and clinical anxiety-related autonomic hyperactivation symptoms, respectively. The convergent autonomic circuit also significantly aligned with causal PTSD and anxiety circuits but not a causal depression circuit, suggesting a specific link to disorders featuring autonomic hyperactivation. Together, these findings support the potential utility of the convergent circuit as a neuromodulation target for autonomic hyperactivation.

Across heterogeneous datasets, the convergent autonomic circuit was robust to differences in outcome measure, stimulation modality, or participant sample (i.e., healthy controls or patients with mild Alzheimer’s disease). These methodological differences introduce noise when searching for convergence, which likely attenuates the observable effect size. Despite this inherent limitation, combining diverse datasets also carries at least two important strengths (34). First, it mitigates study-specific confounders such as spatial bias; TMS targets were selected based on sgACC connectivity, and DBS and LIFU were applied to circumscribed hypothalamic area and insular targets, respectively. Because these biases were unique to each study, convergent circuit effects were unlikely to be driven exclusively by them. Second, it coalesces study-specific strengths; the TMS and LIFU datasets measured single, precisely measured continuous autonomic outcomes, while the DBS dataset measured several categorical autonomic symptoms. The convergent autonomic circuit combines these depth/breadth advantages, linking it more strongly to an underlying autonomic arousal construct.

The convergent autonomic circuit contains several brain areas previously implicated in autonomic arousal (35). For example, the paraventricular hypothalamus, sometimes described as the “master controller” of the autonomic nervous system (36), was a significant negative convergent circuit location. The infundibulum, a hypothalamic output linked to neuroendocrine effects (37), showed the same effect. This effect was not driven solely by the fact that the hypothalamic-area was stimulated in the DBS datasets, as these regions were the top two peaks in the circuit derived only from TMS data (see Supplementary Materials). Convergent autonomic circuit peaks are in the fornix, a fiber tract that terminates in the hypothalamus (26), and the anterior midcingulate cortex (aMCC). Previous neuromodulation results implicate the aMCC in autonomic arousal, whereby direct aMCC electrical stimulation evokes changes in heart rate (28), direct modulation with LIFU alters autonomic responses to pain (29), and subcallosal cingulate DBS evokes changes in heart rate via aMCC structural connections (27). Additional autonomic arousal-relevant regions including the dorsomedial prefrontal cortex (dmPFC), anterior insula, and ventral midbrain (i.e., substantia nigra) were positive connections of the convergent autonomic circuit. The dmPFC is part of a causal anxiety circuit (38) and is thought to contribute to conscious threat appraisal (39). The anterior insula is a key salience network node implicated in the regulation of autonomic activity and awareness of bodily sensations (40). Finally, substantia nigra degeneration is a hallmark of Parkinson’s disease, which features dysautonomia as a core symptom (41).

While derived from brain stimulation sites that induced transient autonomic changes, the convergent circuit was also associated with sustained changes in anxiety-related autonomic activation. In healthy participants, TMS target sites that more strongly intersected the autonomic arousal circuit reduced anxiety-related startle during a laboratory task more than sham 24 hours after stimulation. In Vietnam War Veterans, penetrating brain injury lesions that more strongly intersected the convergent autonomic circuit led to lower levels of PTSD-related exaggerated startle. In patients with moderate to severe depression and anxiety, TMS sites that more strongly intersected the convergent autonomic circuit reduced a cluster of anxiety-related hyperactivation symptoms. The convergent autonomic circuit was also shown to significantly align with causal PTSD and anxiety circuits, but not a causal depression circuit. These findings suggest that the convergent autonomic circuit could potentially serve as a neuromodulation target for autonomic hyperactivation. Alternatively, autonomic outflow might be used to test engagement of existing targets for disorders featuring autonomic hyperactivation (i.e., PTSD and anxiety).

The novelty and strength of this study is robustness across heterogeneous causal datasets that manipulated and measured autonomic arousal differently. However, the study has some key limitations. First, the datasets used to derive the convergent autonomic circuit measured transient changes in autonomic arousal. Encouragingly, the circuit was associated with sustained reductions in anxiety-related autonomic hyperactivation, supporting potential enduring effects of convergent circuit stimulation. However, prospective studies are needed to test this possibility.

Second, the autonomic arousal measures in the present study cannot discriminate between sympathetic and parasympathetic involvement. Future studies should use techniques such as microneurography to more precisely parse sympathetic versus parasympathetic effects of circuit modulation. Third, the included neuromodulation targets did not randomly vary but were localized using functional neuroimaging, or were constrained to specific regions (i.e., hypothalamic area/insula). Although the included lesions were more spatially diffuse, they were also biased to brain areas more likely to be affected by penetrating brain injuries. Importantly, these spatial biases were study-specific, so any consequent biases are mitigated by our finding of convergence across studies. Fourth, a normative connectome was used to map functional connections of brain stimulation and lesion sites. This precluded measurement of individual functional connectivity differences potentially relevant to the present outcome. Future studies that collect sufficient functional connectivity data to generate reliable individual-level maps are necessary to validate the present findings.

In conclusion, the present study mapped a convergent autonomic circuit connecting TMS, DBS, and LIFU sites that transiently reduced autonomic arousal. This circuit was associated with enduring reductions in anxiety-related autonomic activation. Brain stimulation RCTs that prospectively target this circuit and measure its effects on autonomic hyperactivation are warranted.

## Online Methods

### Characteristics of included datasets

We sought to gather datasets that measured the effect of focal brain perturbations (i.e., brain lesions/stimulation) on autonomic arousal. We identified three datasets that measured transient changes in autonomic arousal induced by TMS, DBS, or LIFU applied to different brain areas. These datasets were used to generate distinct autonomic arousal circuits and a convergent autonomic circuit. In post hoc analyses of additional independent datasets, we tested: (1) whether incidental intersection of TMS sites with the convergent autonomic circuit is associated with anxiety-related autonomic activation to a laboratory task; (2) whether the convergent circuit is aligned with causal anxiety-related circuits (14,31), and (3) whether incidental intersection of lesions and TMS sites with the convergent autonomic circuit is associated with sustained changes in anxiety-related autonomic hyperactivation symptoms. Additional description of included datasets can be found in Supplementary Materials.

### Autonomic circuit datasets

The TMS dataset includes 14 healthy participants who received stimulation to 10 different stimulation targets in the prefrontal (5 sites) and parietal (5 sites) cortices (17) (Figure 1a). Targets were manually selected based on individual resting-state functional connectivity to the subgenual anterior cingulate cortex (sgACC). Strongly positive and negative-sgACC connected targets were identified in each hemisphere of the prefrontal and parietal cortex. Two additional targets with near-zero sgACC connectivity were identified in midline prefrontal and parietal locations. Each target was stimulated using a protocol shown to yield temporary heart rate deceleration entrained to TMS. This protocol includes 5-second trains of 10 Hz TMS, which briefly decelerate heart rate, separated by 11-second inter-trial intervals during which heart rate normalizes. The primary outcome was TMS-induced entrainment of heart rate deceleration at each target. One target was removed due to data corruption, leaving 139 total targets.

The DBS dataset includes 58 patients with mild Alzheimer’s disease who received stimulation to a total of 392 sites in the hypothalamic area (20) (Figure 1b). DBS was applied to individual unilateral leads using fixed stimulation parameters (90 μs, 130 Hz, voltage gradually increased in 1.0V increments). As a tolerability measure, the study examined the effects of hypothalamic area DBS on a range of autonomic symptoms due to the hypothalamus’s role in autonomic function. Measured autonomic symptoms included cardiovascular (tachycardia, hypertension), thermoregulatory (warmth, flushing, sweating, coldness), gastrointestinal (nausea), and vestibular (dizziness) changes.

The LIFU dataset includes 23 healthy participants who received stimulation to a total of 46 sites in the anterior and posterior insula (22) (Figure 1c). To measure the effects on insular LIFU on interoceptive and autonomic outcomes, LIFU was applied 200ms before a painful stimulus (i.e., contact heat evoked potential) for 1-second with a 36% duty cycle. The primary autonomic outcome measure was heart rate variability, defined by the standard deviation of normal sinus beats (23).

### Laboratory and clinical anxiety-related autonomic activation datasets

The laboratory dataset includes 28 healthy participants from a randomized, double-blinded, cross-over study. In this experiment, participants received four sessions of active intermittent theta-burst stimulation (iTBS) or sham (order randomized) to right dorsolateral prefrontal cortex targets 24 hours prior to a laboratory anxiety task. Targets were individualized to right dorsolateral prefrontal cortex locations that showed the strongest fMRI activation to high versus low working memory load. This selection procedure was based on the hypothesis that activation of working memory circuitry attenuates threat responses (42). Active iTBS was delivered twice daily (600 pulses consisting of three 50Hz bursts delivered in 200ms intervals for 2s separated by 8s inter-train intervals) after a working memory task, which was intended to prime working memory circuitry. We previously analyzed data from this trial and found that TMS sites that incidentally intersected a previously derived anxiosomatic circuit reduced anxiety potentiated startle (APS) more than sham (43). APS is a measure of the difference in facial electromyography startle response magnitude (i.e., blink response) evoked by white noise delivered at rest versus during anticipation of uncertain electric shock (44). The anxiosomatic circuit contains many of the same brain areas as the convergent autonomic circuit (e.g., medial prefrontal cortex, ventrolateral prefrontal cortex, posterior cingulate) (38). Given this shared circuitry, we sought to test whether incidental intersection with the convergent autonomic circuit would predict TMS effects on anxiety-related autonomic activation.

The PTSD circuit (31) was derived using data from the Vietnam Head Injury Study (VHIS). This dataset includes 196 male Veterans with a penetrating brain trauma lesion who underwent a psychiatric assessment battery. Among other assessments, the battery includes the Clinician Administered PTSD Scale (CAPS) (45) to assess PTSD, the Neurobehavioral Rating Scale (NBRS) (46) to assess neuropsychiatric symptoms including anxiety, and the Beck Depression Inventory (BDI) (47) to assess depression. Three participants in the VHIS dataset did not have PTSD data and were therefore excluded from all analyses. Of the remaining 193 participants, 61 had PTSD, 39 had subthreshold PTSD, and 93 did not have PTSD. The PTSD circuit was generated by mapping the functional connectivity patterns of lesions using a normative connectome and correlating lesion connectivity with PTSD status (controlling for depression levels and lesion size) at each voxel. A depression circuit was also derived from this dataset to test the specificity of the convergent autonomic circuit to anxiety-related hyperactivation. The depression circuit was generated by mapping the functional connectivity patterns of lesions using a normative connectome and correlating lesion connectivity with BDI levels (controlling for lesion size) at each voxel.

The anxiety circuit was derived from a subset of VHIS participants with NBRS anxiety ratings (n = 181), in combination with another brain lesion dataset and two brain stimulation datasets (total N = 562) (14). The other brain lesion dataset includes 270 patients with stroke causing lesions whose anxiety levels were measured using the Hospital Anxiety and Depression Scale (HADS) (48). The two brain stimulation datasets (n = 111) include depression patients who received conventional high-frequency TMS to scalp-based left dlPFC targets. Anxiety was measured using the BDI in one of these datasets and the Hamilton Depression Rating Scale (HAMD) (49) in the other. Unique anxiety circuits were generated in each dataset by mapping the functional connectivity patterns of brain lesions/stimulation sites using a normative connectome and correlating brain lesion/stimulation site connectivity with anxiety levels (controlling for depression levels and lesion size when applicable) at each voxel. These circuits were shown to be more similar than chance and were therefore combined into a convergent anxiety circuit. Additional anxiety data came from a randomized comparative trial (n = 36) that tested the efficacy of a dlPFC target (MNI: -32, 44, 34) against a dmPFC target (MNI: 0, 48, 46) in patients with moderate to severe depression and anxiety. Depression and anxiety were measured using the BDI and BAI, respectively. Six participants who did not have post-treatment symptom measures were removed from the analysis, leaving a total n of 30 (dlPFC: n = 13, dmPFC: n = 17).

### Generating TMS, DBS, and LIFU autonomic circuits

We used causal circuit mapping to generate a TMS circuit associated with greater heart rate deceleration, a DBS circuit associated with less autonomic arousal, and a LIFU circuit associated with greater heart rate variability. First, we estimated stimulation site locations. For TMS, we modeled TMS sites as spheres of decaying intensity (up to 12mm radius) as in prior work (19). We also estimated TMS site locations using electric-field modeling (E-field modeling). However, these results are secondary as E-field modeling has not been validated for use in causal network mapping (see Supplementary Materials). DBS sites were estimated using volume of tissue activation (VTA) modeling (21). LIFU sites were estimated using acoustic modeling (22). After estimating stimulation site locations, we assessed their whole-brain functional connectivity patterns using a normative connectome database generated from healthy participants (12), as in our prior work (13,31,38). Then we employed repeated measures linear mixed-effects models to assess voxel-wise relationships between stimulation site connectivity and autonomic changes for TMS, DBS, and LIFU separately. In these models, stimulation site connectivity (predictor variable) was regressed on autonomic changes (outcome variable), controlling for within-subject effects. This yielded a t-value at each voxel corresponding to the strength of the relationship between stimulation site connectivity and autonomic changes.

Positive t-values reflect positive connections of brain stimulation sites that reduce autonomic arousal; negative t-values reflect the opposite.

For TMS, we assessed the relationship between TMS site connectivity and heart rate deceleration. For LIFU, we assessed the relationship between LIFU site connectivity and heart rate variability. For DBS, we assessed relationships between DBS connectivity and a broad range of DBS-induced autonomic arousal symptoms including tachycardia, hypertension, warmth, flushing, sweating, coldness, nausea, and dizziness. We combined symptoms from the same category to prevent overweighting of specific categories. Tachycardia and hypertension were combined into a cardiovascular category; warmth, flushing, sweating, and coldness were combined into a thermoregulation category; nausea formed a single-item gastrointestinal category; and dizziness formed a single-item vestibular category. Categories were given overall scores of 0 or 1; participants with no symptoms within a category received a 0, while participants with one or more symptoms received a score of 1. Autonomic arousal symptom scores equaled the sum-total of cardiovascular, temperature, gastrointestinal, and vestibular symptoms. In secondary analyses, we analyzed autonomic arousal in different ways to ensure that our results were not biased by analytical choices. This involved summing individual autonomic arousal symptoms (rather than summing across categories), generating circuits from individual symptoms and then averaging them, and generating averaged circuits after leaving one symptom out.

### Generating a convergent autonomic circuit

We tested for convergence between TMS, DBS, and LIFU maps using Pearson correlation. Specifically, we correlated between each pair and Fisher z-transformed the results before averaging them to generate a pooled correlation value. We tested the significance of this result using a permutation method that was similar to our previous work (13,38) but that maintained the within-subject structure of each dataset. For the TMS and DBS datasets, we randomly permuted clinical outcomes associated with a given stimulation site with a different stimulation site for the same subject. For the TFUS dataset, this approach was not optimal because each subject only had two stimulation targets. Therefore, we performed a block permutation in which clinical outcomes from both targets were randomly assigned together to another subject. The resulting permuted TMS, DBS, and LIFU maps were correlated with one another as described above to create a null distribution against which the real correlation was compared. The two-tailed significance threshold was set at p < 0.05.

After the TMS, DBS, and LIFU circuits were shown to be more similar than expected by chance, they were combined into a convergent autonomic circuit. This involved converting circuit values from t-values to r-values, Fisher z-transforming r-values, calculating a mean z’ circuit weighted by sample-size, transforming z’-values back to r-values, and then converting r-values back to t-values.

Statistical significance of the convergent autonomic map was determined using Westfall-Young permutation methods, following the recommendations of Winkler et al. (2014) (25). The same shuffling method described above was employed to preserve the within-subject structure of the dataset. A total of 1,000 permutations were performed, creating a null distribution against which real results were compared. Voxels with t-values greater than 95% of null values were considered significant.

### Assessing relevance to laboratory and clinical anxiety-related activation

We tested the convergent autonomic circuit’s relevance to anxiety-related activation in a series of post hoc analyses. To test whether TMS site functional connectivity to the convergent autonomic circuit predicts laboratory activation, we estimated TMS site locations modeled using the same method described above. We calculated intersection with the convergent autonomic circuit by estimating functional connectivity of each TMS site using the same normative connectome and correlating the resulting seed-maps with the convergent autonomic circuit. We Fisher z-transformed the correlation values and entered them into linear regression models that tested whether connectivity to the convergent autonomic circuit could predict laboratory anxious arousal measured using anxiety potentiated startle.

We also performed a series of post hoc analyses examining relationships between the convergent autonomic circuit and clinical anxiety-related hyperactivation. We performed spatial Pearson correlation to test alignment with the previously described causal PTSD, anxiety, and depression circuits. Significance was determined using the same permutation method employed in the TMS, DBS, and TFUS correlation analysis. One-tailed testing was used because in our prior work we proposed that stimulation of the medial prefrontal cortex would reduce symptoms of PTSD and anxiety (14,31), disorders characterized by autonomic hyperactivity. Therefore, a result in the opposite direction would be considered a failed test for that hypothesis. We also tested whether functional connectivity of combat brain lesions to the convergent autonomic circuit is associated with lifetime PTSD symptom levels measured via the CAPS. Lesions were traced using computed tomography scans and converted to MNI space (for additional detail, see: 49). Their functional connectivity patterns were then mapped using the same normative connectome and correlated with the convergent autonomic circuit. Spearman correlation was used to test relationships between Fisher z-transformed functional connectivity results and PTSD symptom levels, controlling for depression symptoms and lesion size (31). We also tested whether functional connectivity of TMS sites to the convergent autonomic circuit predicts changes in anxiety-related autonomic hyperactivation symptoms measured using the BAI. Hyperactivation symptoms included: numbness or tingling, feeling hot, wobbliness in legs, dizzy or lightheaded, heart pounding/racing, unsteady, feeling of choking, hands trembling, shaky/unsteady, difficulty in breathing, indigestion, faint/lightheaded, face flushed, and hot/cold sweats. Participants were randomized to one of two standardized MNI target locations, limiting incidental variability in target location. However, high-quality individual fMRI data were collected, enabling reliable measurement of individual differences in functional connectivity to each target. Therefore, we generated individualized target seed-maps using each participant’s resting-state fMRI data. We then correlated these seed-maps with the convergent autonomic circuit, Fisher z-transformed the result and correlated it with changes in hyperactivation symptoms separately for each target condition, controlling for baseline hyperactivation, baseline depression, and changes in depression. We performed separate analyses for each condition because equal variance between targets could not be assumed. We combined results by Fisher z-transforming Spearman r correlation values, generating a weighted z-value, and calculating a pooled p-value from the weighted z-value and combined standard error.

## Supplementary Materials

**Supplementary Table 1:** The convergent autonomic circuit was derived from three datasets that used different neuromodulation methods and measured transient changes in different autonomic arousal outcomes. The circuit was validated using additional independent brain stimulation and lesion datasets that measured sustained changes in anxiety-related autonomic activation outcomes.

|  | Modality | n | Patient population | Primary outcome | Age mean, sd | Sex |
| --- | --- | --- | --- | --- | --- | --- |
| Autonomic circuit datasets | TMS | 14 (139 stimulation sites) | Healthy participants | Heart rate deceleration as measured using heart-brain coupling | 36.4 (14.7) | 60% M, 40% F |
|  | DBS | 58 (392 stimulation sites) | Mild Alzheimer's disease | Multiple categorical autonomic symptoms | 68.5 (7.9) | 55% M, 45% F |
|  | TFUS | 23 (46 stimulation sites) | Healthy participants | Heart rate variability as measured using standard deviation of normal sinus beats | 27 (5.5) | 30% M, 70% F |
| Anxiety-related autonomic activation datasets | TMS | 28 | Healthy participants | Anxiety potentiated startle | 23.75 (4.3) | 36% M, 64% F |
|  | Lesions | 193 | Veterans with penetrating brain injuries | Clinician Administered PTSD Scale | 58.3 (3.1) | 100% M |
|  | TMS and brain lesions | 562 | TMS: patients with depression<br>Lesions: patients with penetrating brain injuries or ischemic/hemorrhagic strokes | TMS: Beck Depression Inventory or Hamilton Depression Rating Scale symptom clusters<br>Lesions: Neurobehavioral Rating Scale Anxiety Item, Hospital Anxiety and Depression Scale | TMS: 53 (10); 47 (11)<br>Lesions: 58 (3); 66 (12) | TMS: 33% M, 67% F; 43% M, 57% F<br>Lesions: 100% M; 61% M, 39% F |
|  | TMS | 30 | Patients with depression and anxiety | Beck Anxiety Inventory | 34.36 (11.81) | 47% F, 53% M |

**Supplementary Figure 1:**
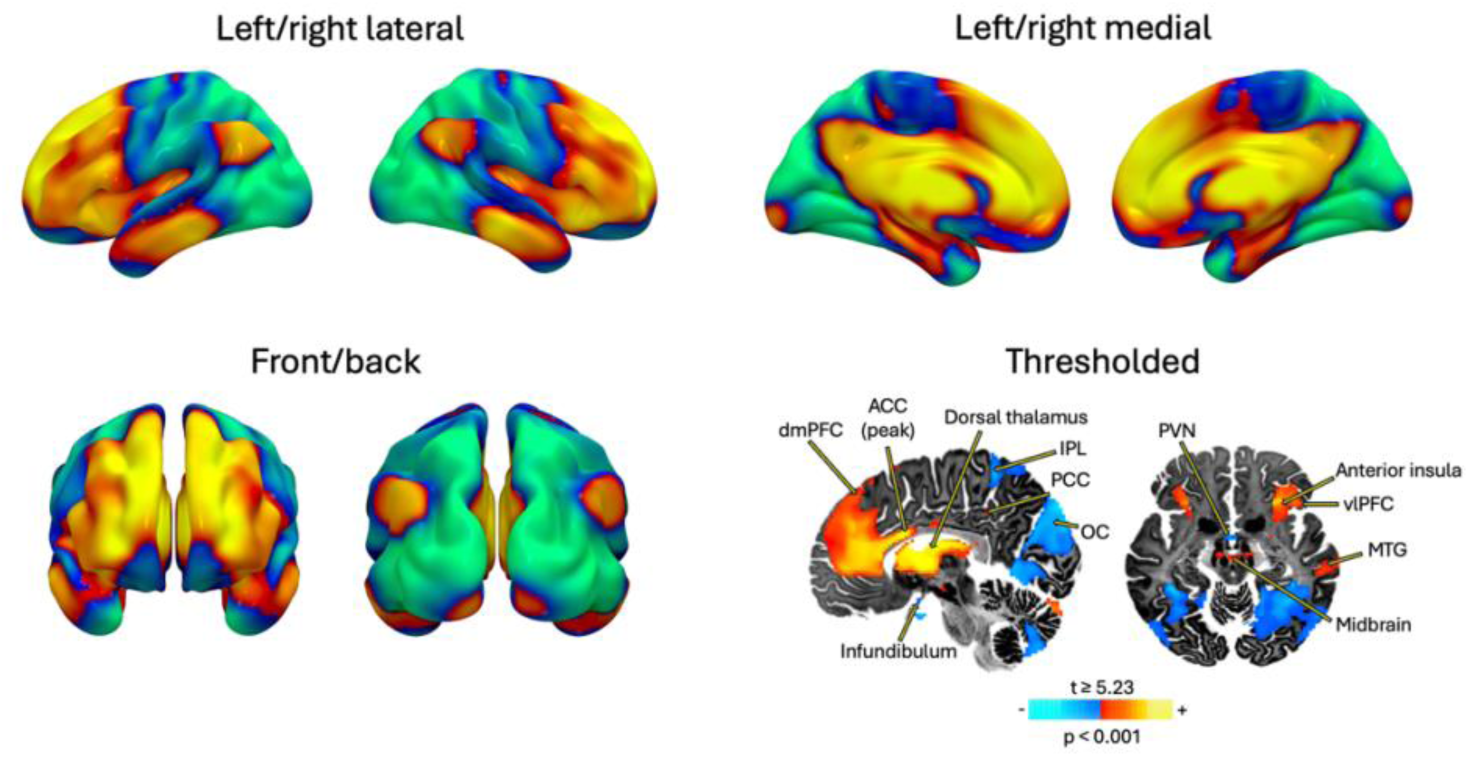
Convergent autonomic circuit

**Supplementary Figure 2:**
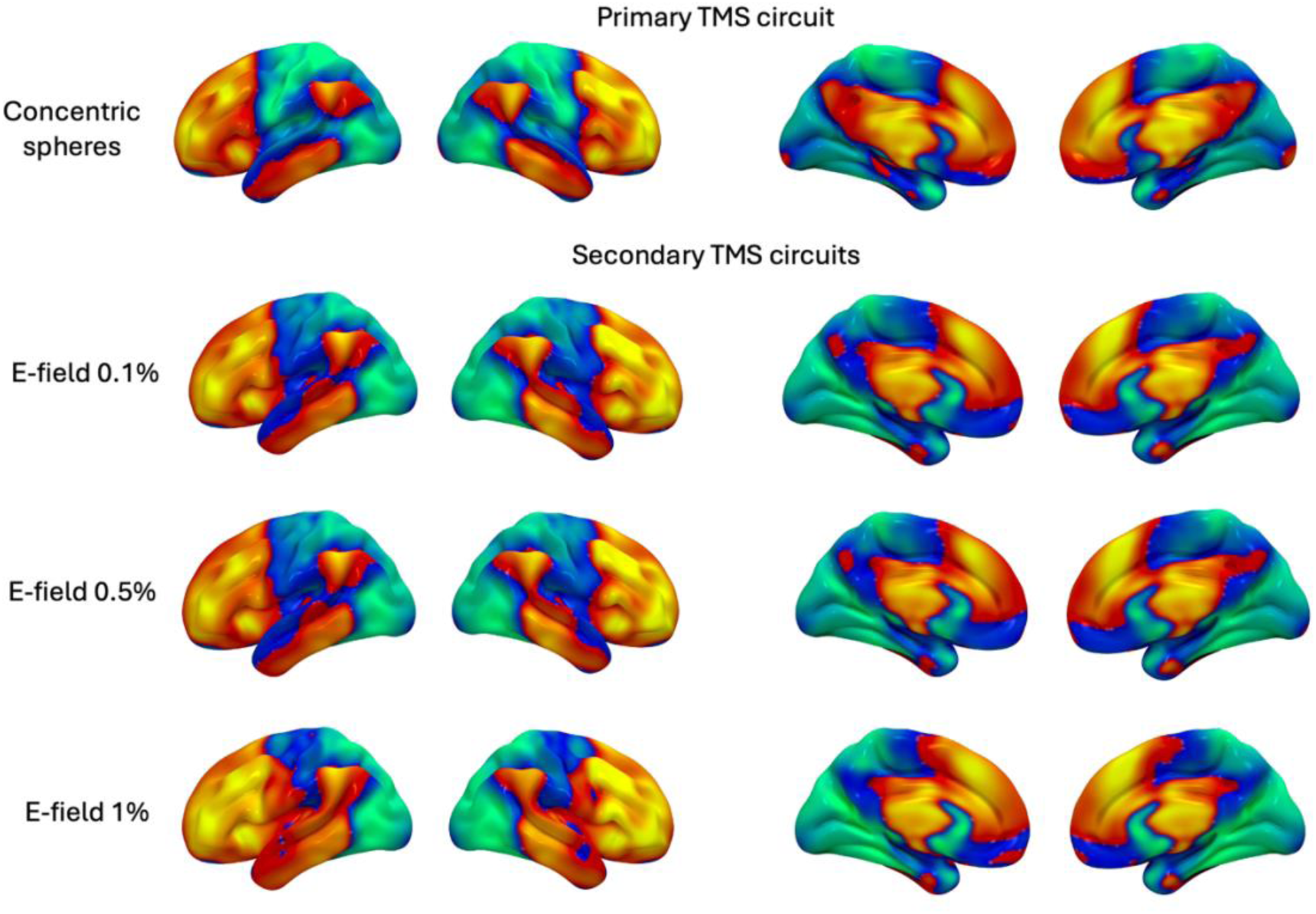
TMS circuit variations

**Supplementary Figure 3:**
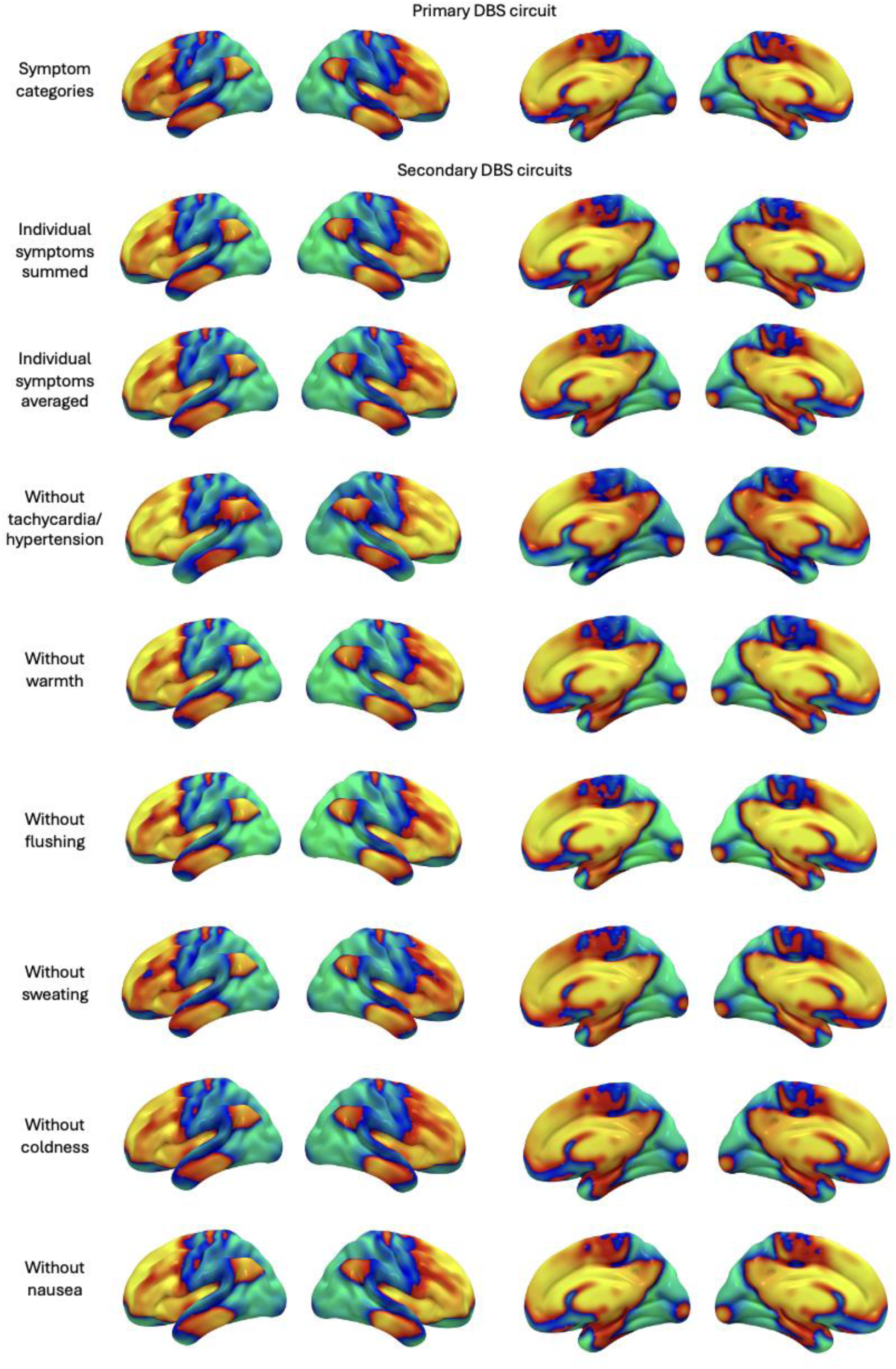
DBS circuit variations

**Supplementary Figure 4:**
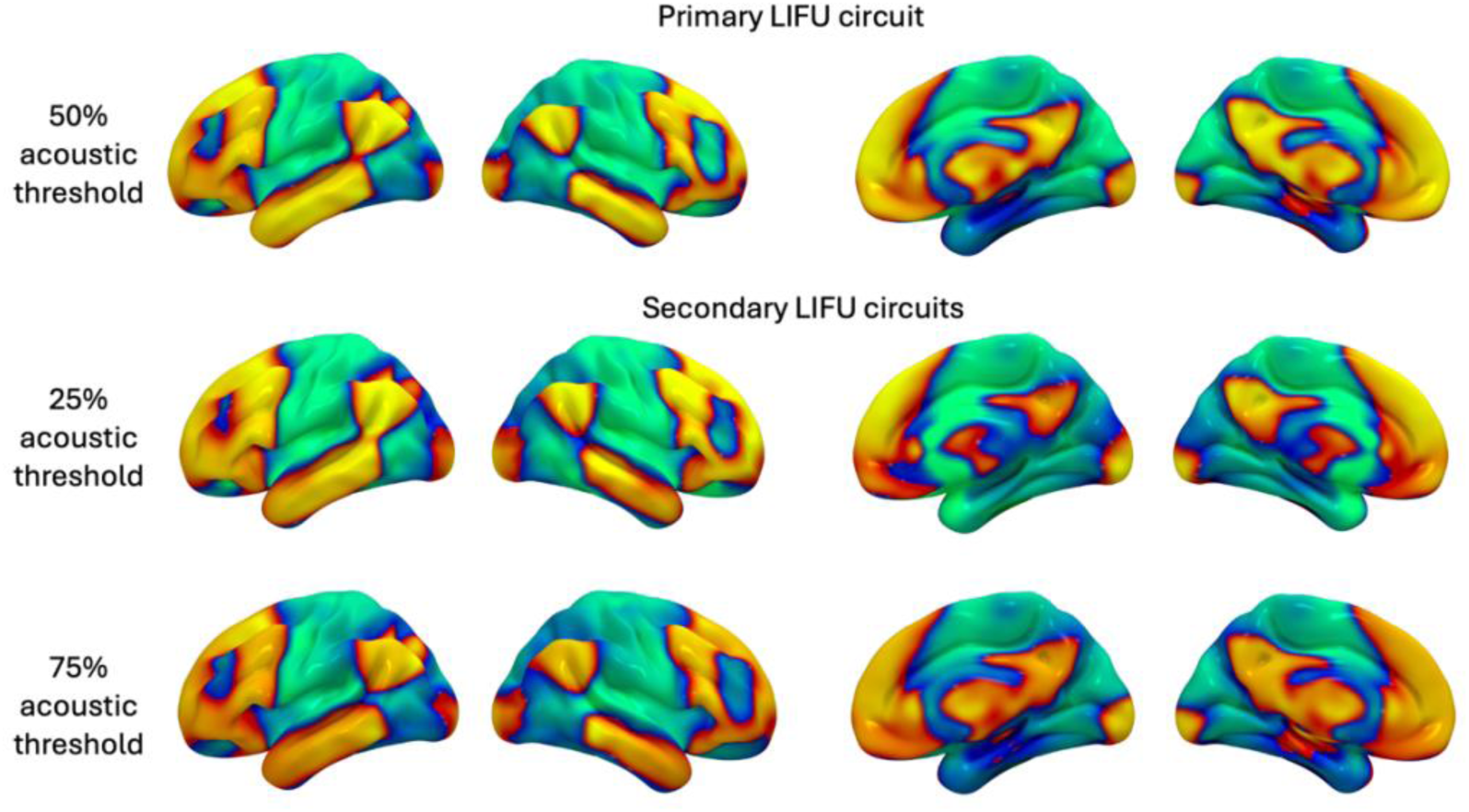
LIFU circuit variations

**Supplementary Figure 5:**
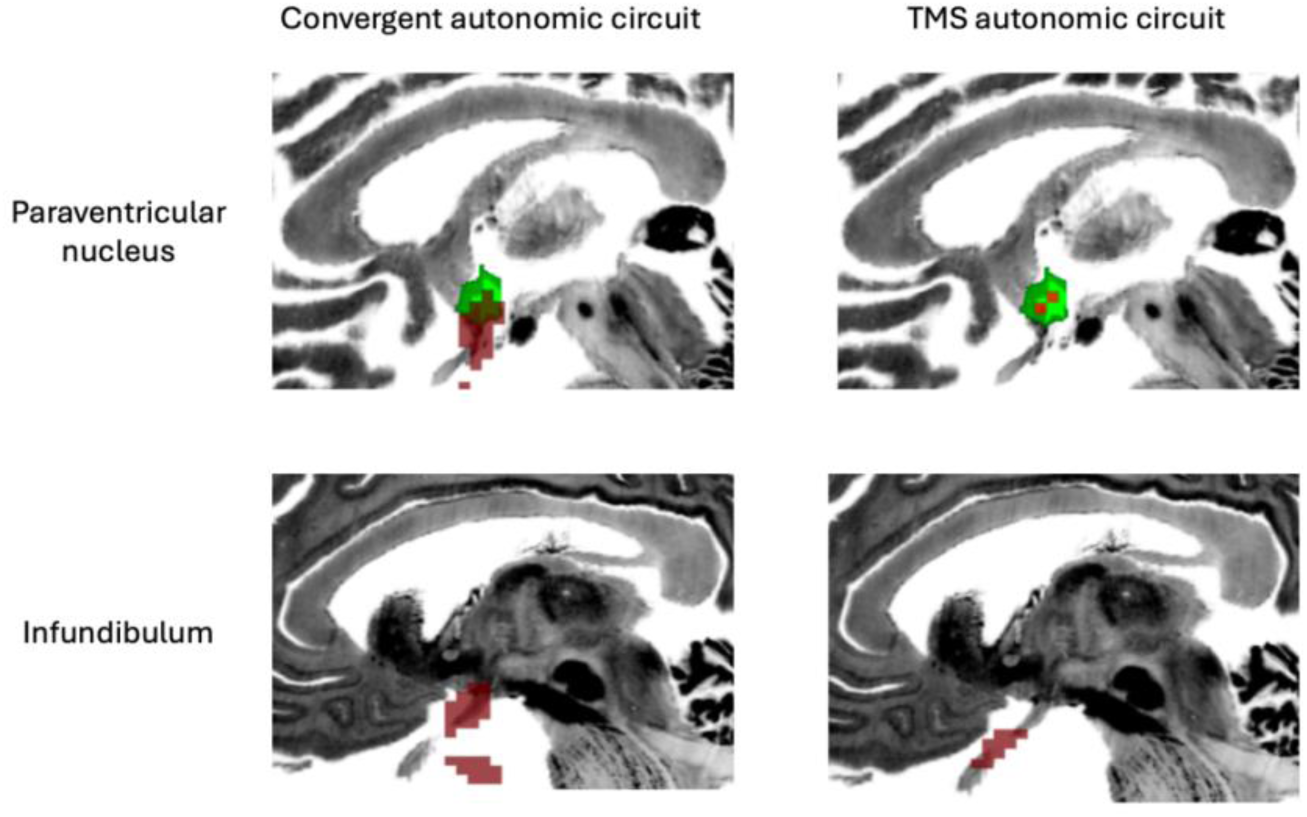
The paraventricular nucleus of the hypothalamus is depicted in green. Significant convergent autonomic circuit (top left pane) and TMS autonomic circuit (top right pane) paraventricular nucleus locations are depicted in red. Significant infundibulum circuit locations are shown in the bottom panes.

## Notes

### Competing Interest Statement

The authors have declared no competing interest.

